# Establishment of the Meyer-Overton correlation in an artificial membrane without protein

**DOI:** 10.1101/2023.10.18.562855

**Authors:** Atsushi Matsumoto, Yukifumi Uesono

## Abstract

**Background:** The potency of anesthetics with various structures increases exponentially with lipophilicity, which is the Meyer-Overton (MO) correlation discovered over 120 years ago. The MO correlation was also observed with various biological effects and various chemicals, including alcohols; thus, the correlation represents a fundamental relationship between chemicals and organisms. The MO correlation was explained by the lipid and protein theories, although the debate has not been resolved and the principle remains unknown.

**Methods:** The gentle hydration method was used to form giant unilamellar vesicles (GUVs) consisting of high- and low-melting phospholipids and cholesterol in the presence of *n*-alcohols (C_2_-C_12_). Confocal fluorescence microscopy was used to determine the percentage of GUVs with domains in relation to the concentrations of *n*-alcohols.

**Results:** *n*-Alcohols inhibited the domain formation of GUVs, and the half inhibitory concentration (IC_50_) in the aqueous phase decreased exponentially with increasing chain length (lipophilicity). In contrast, the membrane concentrations of alcohols for the inhibition, which is a product of the membrane-water partition coefficient and the IC_50_ values, remained constant irrespective of the chain length.

**Conclusions:** The MO correlation is established in artificial lipid membrane, which supports the lipid theory. When alcohols reach the same critical concentration in the membrane, similar biological effects appear irrespective of the chain length, which is the principle underlying the MO correlation.

**Summary Statement:** *n*-Alcohols inhibit the domain formation of giant unilamellar vesicles according to the Meyer-Overton correlation.

## Introduction

The potency of anesthetics with various structures increases exponentially with lipophilicity, which is the Meyer-Overton (MO) correlation discovered over 120 years ago.^1^ The MO correlation was observed with various biological effects by various chemicals, such as alcohols, alkanes, fatty acids, antipsychotic drugs, and agrochemicals, in addition to anesthetics.^2–8^ Therefore, the correlation represents a fundamental relationship between chemicals and organisms and has led to the notion of a quantitative structure−activity relationship (QSAR); this relationship is used to estimate the biological effects of various chemicals from their calculated lipophilicities *in silico*,^8^ although the principle remains unknown. On the basis of the MO correlation, the cell membrane was determined to be lipophilic, which is an important concept. As a result, the lipid theory was proposed, which states that chemicals dissolve in the lipophilic membrane and act on various membrane targets indirectly.^9^ However, the theory failed to explain a cutoff paradox in which long-chain alcohols (≥C_13_) lose their biological effects, as these compounds should exhibit strong effects according to the MO correlation.^4,10^ Instead, the protein theory was proposed based on luciferase inhibition by anesthetics and alcohols in the absence of a lipid membrane. The theory advocated that the chemicals directly bind proteins and explained the cutoff by assuming that ideal pockets accommodate various chemicals but not the long-chain alcohols.^11^ However, the pocket has not been revealed to date. These two hypotheses have been debated for a long time because elucidating the common mechanisms underlying the action of numerous chemicals, including anesthetics, that obey the MO correlation is crucial. The conclusions obtained from debate should not be ambiguous because the knowledge will determine the fundamental direction for drug design based on whether the main axis is the interaction with lipids or proteins.

We previously demonstrated that the cutoff of long-chain alcohols resulted from the low concentration of effective monomer hydrated with water, and the alcohols did not reach the critical concentration necessary to exert biological effects.^12^ Thus, when the cutoff alcohols are sufficiently accumulated into the membrane, strong biological activity is observed according to the MO correlation, indicating that the lipid theory can reasonably explain the correlation.^13^ Reproducing the MO correlation in the lipid membrane free from proteins could provide more direct evidence to support the lipid theory, but this evidence has not been attained thus far. Recently, general anesthetics were reported to disrupt lipid rafts in living cells.^14^ However, it is difficult to distinguish whether the disruption results from the interaction between anesthetics and lipids or proteins. This is because lipid rafts consisting of lipids and proteins are highly dynamic and unstable nanodomains that associate/dissociate rapidly via protein‒protein and protein-lipid interactions.^15^ In contrast, the lipid domains formed in artificial membranes are stable because there are no proteins. Here, we demonstrated the MO correlation in the artificial membrane by monitoring alcohol-induced inhibition of membrane domain formation.

## Materials and Methods

### Chemicals

Phospholipids, 1,2-distearoyl-*sn*-glycero-3-phosphocholine (18:0-PC, DSPC), 1,2-dioleoyl-*sn*-glycero-3-phosphocholine (18:1-PC, DOPC), 1,2-distearoyl-*sn*-glycero-3-[phosphor-*rac*-(1-glycerol)] (18:0-PG, DSPG), and 1,2-dioleoyl-*sn*-glycero-3-[phosphor-*rac*-(1-glycerol)] (18:1-PG, DOPG) were purchased from Avanti Polar Lipids (Alabaster, AL). Cholesterol (CHOL) and *n-* alcohols (C_2_-C_10_) were obtained from Wako Pure Chemical (Tokyo Japan). The fluorescent dye 1,1’-didodecyl-3,3,3’,3’-tetramethylindocarbocyanine perchlorate (C12:0-DiI) was from Thermo Fisher Scientific (Waltham, MA). Naphtho-[2,3a]-pyrene and 1-dodecanol (naphthopyrene) were obtained from Tokyo Chemical Industry (Tokyo, Japan). All materials were used without further purification. The phospholipid concentration was determined using the inorganic phosphate assay.^16^ CHOL and alcohol concentrations were determined analytically. The fluorescent dye concentration was determined using a UV‒Vis spectrophotometer (U-3010, Hitachi, Tokyo, Japan). Extinction coefficients in MeOH were 143,000 cm^-1^M^-1^ at 549 nm for C12:0-DiI according to the manufacturer (Thermo Fisher Scientific) and 23,749 cm^-1^M^-1^ at 454 nm for naphthopyrene.^17^

### Preparation of Giant Unilamellar Vesicles (GUVs)

GUVs were prepared by a gentle hydration method as described previously^18,19^ with modifications for the use of alcohols. Ten mol% of DSPC and DOPC were substituted by DSPG and DOPG, respectively, because repulsion between negatively charged head groups is necessary to obtain GUVs by this method. The presence of PGs does not affect the phase behavior in the absence of alcohols.^18^ Lipid mixtures (250 nmol) of the indicated composition with 0.1 mol% naphthopyrene and 0.02 mol% C12:0-DiI were dissolved with 100 μL chloroform in a test tube and dried into a thin film on a rotary evaporator at 60 °C, followed by 1 h with high vacuum. The thin dry film was then hydrated by introducing wet N_2_ gas at 60 °C for 25 min. Two mL of prewarmed 100 mM glucose solution was added to the indicated concentration of the alcohols and incubated for 2 h at 60 °C for GUV formation, followed by gently cooling over 10 h to 25 °C. The GUVs were harvested into 100 mM sucrose solution containing the same concentration of alcohols so that glucose-containing GUVs floated in the sucrose solution. The scheme is shown in Supplemental Figure S1.

### Microscopic Analysis

GUVs were mounted on a slide using a silicon spacer (0.25 mm) placed between the slide and a coverslip. An LSM 510 META microscope (Carl Zeiss, Tokyo, Japan) with a 488/543 nm beam splitter filter and a 40× 1.3 NA oil-immersion objective was used for confocal fluorescence microscopy. Naphthopyrene was observed at 458 nm excitation and 500/520 IR emission, C12:0-DiI at 543 nm excitation and 560 nm longpass emission. The 488/543 nm filter was not ideal for 458 nm excitation, but clear complementary partitioning of the two dyes was observed. The sample slide was observed at room temperature (25-30℃) or at a temperature controlled with a laboratory-made temperature stage consisting of a Peltier device, a water circulating heat sink and a thermomodule controller (MT863-04C06 PLUS, Netsu Denshi Kogyo, Tokyo, Japan). The unilamellarity of GUVs was judged by a previous method^19^, and the vesicles with the lowest brightness were used for analysis. Images were analyzed using the ZEN 3.0 blue edition (Carl Zeiss) and the Fiji distribution of ImageJ 1.53c.^20^ Every optical slice with a 2 mm increment was observed over whole GUVs, and clear complemental localization of C12:0-DiI and naphthopyrene was judged as the presence of membrane domains.

### Data Analysis and Statistics

The 50% inhibitory concentration (IC_50_) of GUV formation with domains was determined by the nonlinear least-squares method^21^ of the log-logistic models^22^ using Solver add-in bundled with Microsoft Excel (Microsoft Corporation, Redmond, WA). Statistical analyses were performed by one-way analysis of variance with Dunnett’s multiple comparison test using the R statistical software package ver. 4.0.3 (The R Foundation for Statistical Computing, Vienna, Austria).

## Results

### Formation of various domain structures in giant unilamellar vesicles

Some proteins and lipids segregate into distinct membrane domains at the nanoscale to microscale. The membrane domains are conserved in animals, fungi, plants, and prokaryotes^23,24^ and participate in various cellular processes, such as signal transduction.^25^ Alcohols are known to affect the phase transition temperature of artificial membranes free from proteins,^26^ although it is difficult to realize the biological significance for scientists that investigate drug effects at a constant temperature. The phase transition temperature was lowered in alcohol-treated giant plasma membrane vesicles (GPMVs) prepared from rat basophilic leukemia (RBL-2H3) cells. The phase transition was visualized as a change to a single liquid domain from the liquid-ordered (L_o_) and liquid-disordered (L_d_) coexisting domain at the same temperature.^27^ Because the lipid-lipid interaction is important for domain formation,^28^ alcohols might inhibit the interaction. However, in GPMVs containing proteins and lipids, it is difficult to distinguish whether alcohols inhibit lipid interactions directly or indirectly via proteins. Similar domains are formed in the artificial lipid membrane and readily observed in giant unilamellar vesicles (GUVs) with diameters greater than 10 μm through fluorescence microscopy.^29^ For instance, in GUVs consisting of high- and low-melting phospholipids and cholesterol (CHOL), highly packed L_o_ domains and L_d_ domains have been observed,^30^ although there is no report of MO correlation by alcohols in GUVs.

We constructed representative GUVs consisting of DSPC, DOPC, and CHOL.^18^ To mimic GPMVs, 10 mol% DSPC and DOPC were substituted by negatively charged lipids, DSPG and DOPG, respectively. The domains were stained with the fluorescent dyes C12:0-DiI (red) for the L_d_ domains and naphthopyrene (blue) for the L_o_ domains (fig. 1C), as reported previously.^18,31^ Various GUVs with different lipid compositions (DSPC/DOPC/CHOL) are summarized in the triangular phase diagram (Gibbs diagram; fig. 1A). Round shapes of the L_d_ domains (fig. 1C) were found near the left side in the boundary of L_d_ + L_o_ coexistence, whereas those of the L_o_ domains (data not shown) were found near the right side (fig. 1A, green-filled circles). These observations indicate that both domains exhibit liquid properties and that this liquid‒liquid coexistence is analogous to the domains observed in GPMVs.^32^ Irregular shapes (fig. 1D) were observed in the low cholesterol region (fig. 1A, red squares). Because C12:0-DiI (red) are enriched in the L_d_ domains,^31^ the domains with enriched naphthopyrene (blue) are the solid gel (L_β_) domains. In contrast, these domain structures disappeared in the high cholesterol region (fig. 1A, white circles). Thus, domain formation with these shapes is largely affected by the amphiphilic cholesterol content. These results were consistent with a previous report with different dyes.^18^

**Fig. 1.**
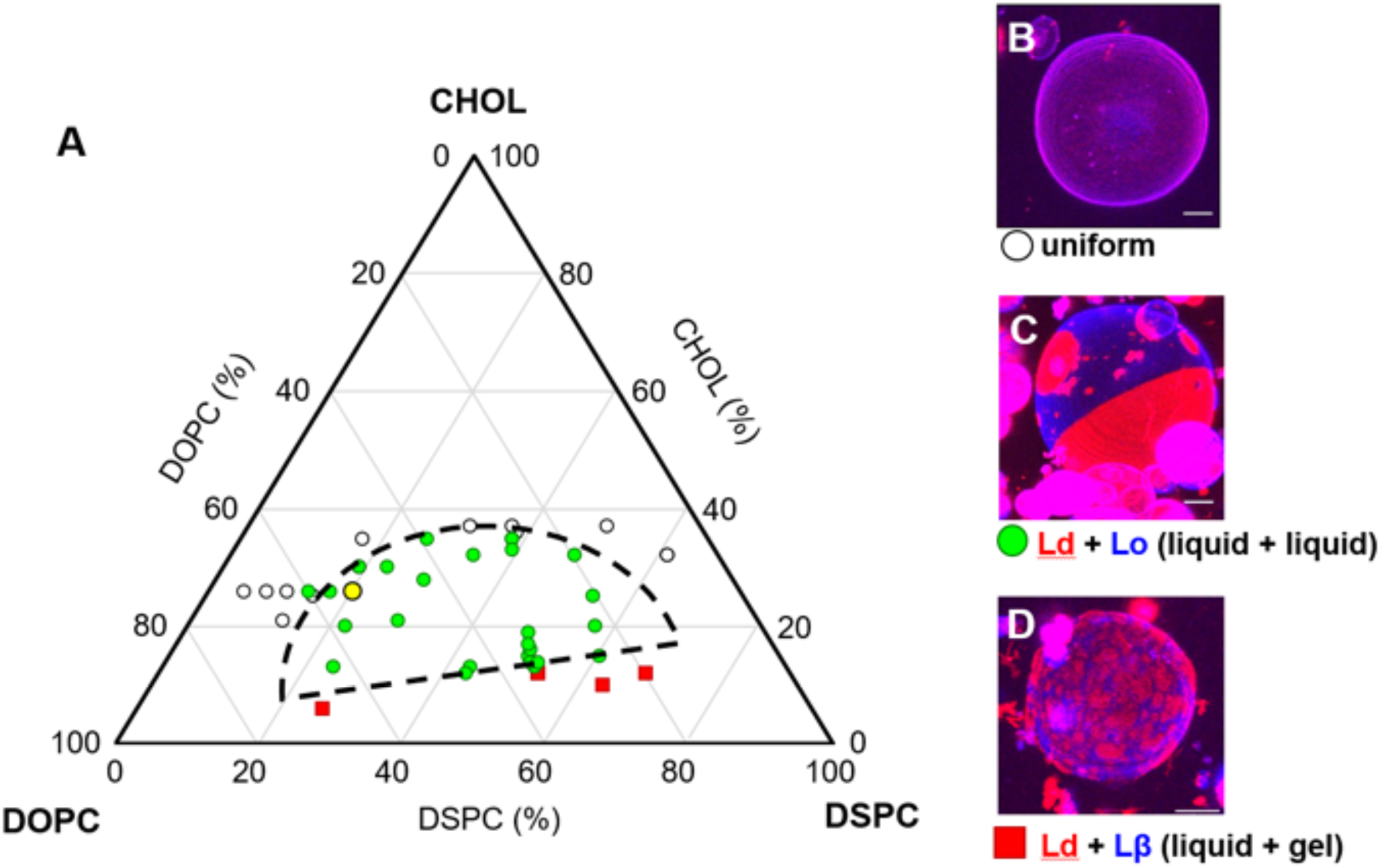
The domain structures of GUVs consisting of different lipid composition. (*A*) A Gibbs diagram for the relationship between the lipid composition of GUVs and the domains observed. (*B*) Open circles indicate the GUV structure without domains. (*C*) Green circles indicate the structure with round-shaped domains. (*D*) Red squares indicate the structure with the non-round shaped domains. The liquid-disordered (L_d_) domains are rich in C12:0-DiI (red), but the liquid-ordered (L_o_) and solid gel (L_β_) domains are rich in naphthopyrene (blue).^31^ The dotted line indicates the assumed boundary of L_d_ + L_o_ region. The yellow circle indicates the GUVs with round domains consisting of lipid composition (DSPC/DOPC/CHOL = 20/54/26). Images were recorded by confocal microscopy z-scans in 2-μm increments. Scales bars are 10 μm.

### Ethanol inhibits the domain formation of GUVs in the absence of proteins

The domains of GPMVs are not formed at high temperatures >25℃.^33^ A similar temperature-sensitive domain is also formed in the GUVs near the upper-left boundary region in which the L_d_+L_o_ domains coexist.^34^ Thus, we examined the effect of alcohols on the GUVs with GPMV-like domains (DSPC/DOPC/CHOL = 20/54/26, fig. 1A, yellow circle). Ethanol did not disappear the preformed domains of GUVs at even 5 M within 15 min (Supplemental Figure S2), whereas it disappeared those of the GPMV at 120 mM within 15 min.^27^ When the GUVs were prepared in the presence of ethanol (C_2_), the percentage of GUVs with domains decreased in a concentration-dependent manner (fig. 2). This result suggests that ethanol affects in a process of domain formation in GUVs. According to the dose‒response curves, the half-inhibitory concentration (IC_50_) of the domain formation was determined to be 1.5 M (fig. 2A). This IC_50_ value was lower than the minimum growth inhibitory concentration (MIC: 2.5 M) (Table 1)^12^ and half lethal dose (LD_50_: 2.3 M) for yeast,^13^ suggesting that domain formation is inhibited at concentrations lower than those exerting biological effects. At 5 M ethanol, almost no domains were formed, although the vesicle structures were maintained. These results suggest that ethanol exerts biological activity by inhibiting membrane domain formation via direct interaction with lipids without destroying the lipid bilayer structure.

**Fig. 2.**
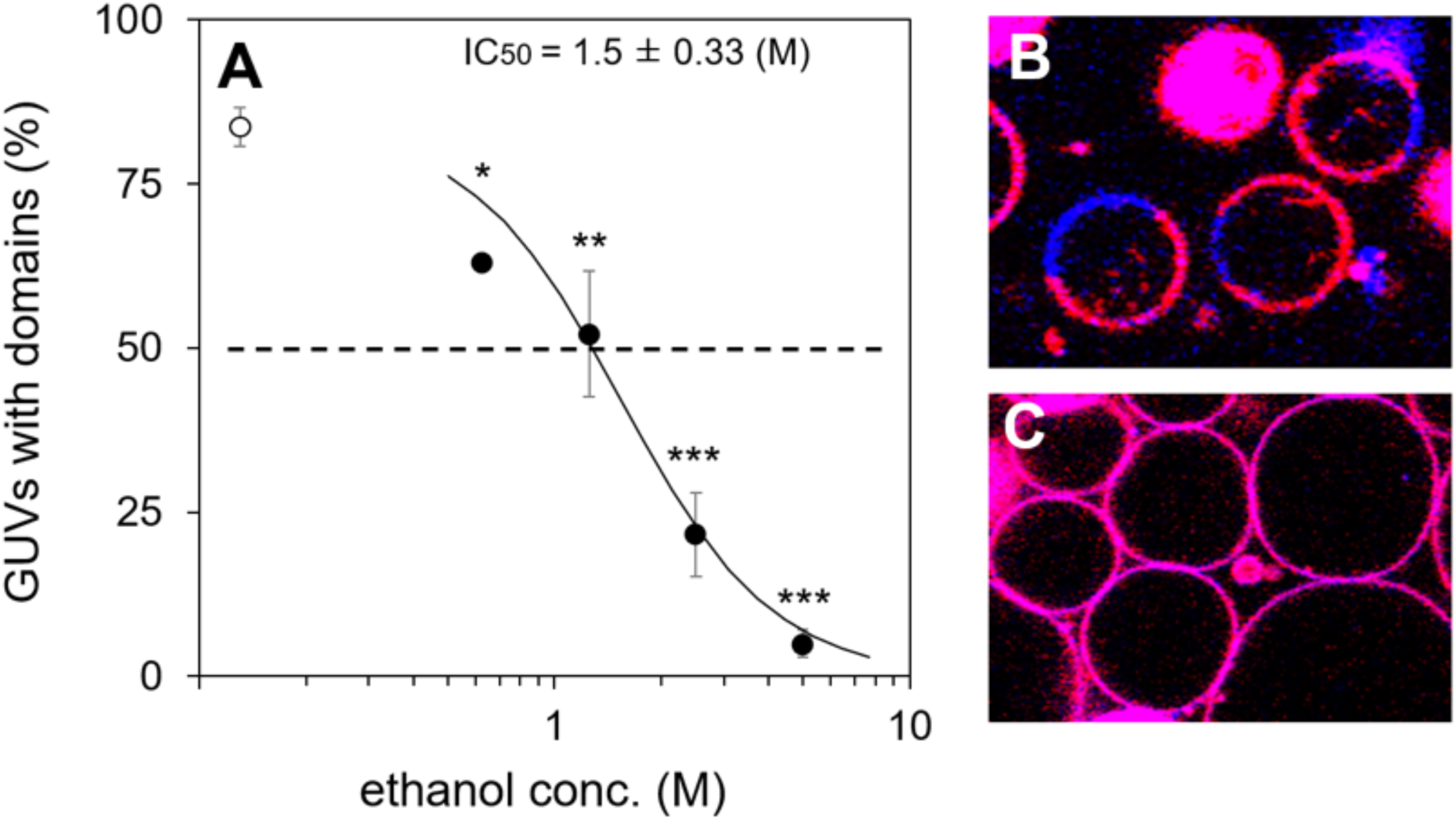
The effect of ethanol on the domain formation of GUVs. (*A*) The percentages of the GUVs with domains are plotted against ethanol concentration (filled circles). An open circle is control without ethanol. The lipid composition of GUV was the same as that in the yellow circle in fig. 1. Images were recorded by confocal microscopy and more than 100 GUVs were counted for each ethanol concentration in one experiment. The logistic curve is fitted to determine 50% inhibitory concentration (IC_50_) and the averages are indicated by lines. Data are the means ± SEM (*n* = 3). Confocal images of GUVs formed in the absence of ethanol (*B*) and in the presence of 5 M ethanol (*C*). Stars indicate significant difference from ctrl (*, *p* < 0.05; **, *p* < 0.01; ***, *p* < 0.001; one-sided Dunnett’s test).

**Table 1.** Physicochemical properties and biological concentrations of alcohols.

| Chain length | $P_{ow}^a$ | $K_{mw}^b$ | $IC_{50}^c$ | $MIC^d$ | $P_{ow} \times IC_{50}$ | $K_{mw} \times IC_{50}$ | $P_{ow} \times MIC$ | $K_{mw} \times MIC$ |
| --- | --- | --- | --- | --- | --- | --- | --- | --- |
| | $^eC_o/C_w$ | $C_m/C_w$ | $C_w$ | $C_w$ | $C_o$ | $C_m$ | $C_o$ | $C_m$ |
| 2 | 0.49 | 0.14 | $1500 \pm 330$ | 2500 | 735 | 210 | 1225 | 350 |
| 4 | 7.6 | 1.5 | $120 \pm 23$ | 250 | 910 | 180 | 1900 | 375 |
| 6 | 107 | 13 | $6.1 \pm 0.70$ | 20 | 653 | 79 | 2140 | 260 |
| 8 | 1000 | 152 | $0.50 \pm 0.039$ | 2.5 | 500 | 76 | 2500 | 380 |
| 10 | 37154 | 1222 | $0.097 \pm 0.032$ | 0.25 | 3604 | 119 | 9289 | 306 |
| 12 | 134896 | 5365 | $0.024 \pm 0.0025$ | 0.063 | 3238 | 129 | 8498 | 338 |
| Max/Min<sup>f</sup> | 275423 | 38321 | 62500 | 39683 | 7.2 | 2.8 | 7.6 | 1.5 |

<sup>a</sup>Octanol/water partition coefficient from PubChem (https://pubchem.ncbi.nlm.nih.gov/).
<sup>b</sup>Membrane/water partition coefficient.<sup>4</sup> <sup>c</sup>Half inhibitory concentration for membrane domain formation determined in fig. 3. Data are the means ± SEMs ( $n = 3$ ). <sup>d</sup>Minimum inhibitory concentration for yeast growth.<sup>12</sup> <sup>e</sup>Concentrations ( $C$ : mM) in different phases are expressed as $C_o$ (in octanol), $C_w$ (in water), and $C_m$ (in membrane). <sup>f</sup>Ratios of the maximum and minimum values in each column ( $C_2$ - $C_{12}$ ).

### *n*-Alcohols inhibit the domain formation of GUVs according to the MO correlation

*n*-Alcohols are known to significantly lower the transition temperature from the L_o_/L_d_ coexisting phases to a single liquid phase of GPMVs with increasing chain length.^27^ Additionally, in GUVs, alcohols (C_4_-C_12_) inhibited domain formation in a dose-dependent manner (fig. 3A). Their IC_50_ values decreased exponentially with increasing chain length (C_2_-C_12_), indicating that the MO correlation is established in the lipid membrane alone. (fig. 3B). C_6_ or C_8_ did not inhibit domain formation efficiently at high concentrations (fig. 3A). This phenomenon was observed in the inhibition of yeast actin polarization at high concentrations of various amphiphiles, such as local anesthetics, quaternary ammonium compounds, and alcohols.^6^ Thus, the inhibition of domain formation in GUVs by alcohols is not specific and might result from the physical property of the amphiphilic structure common to these compounds, although the mechanism is unclear.

**Fig. 3.**
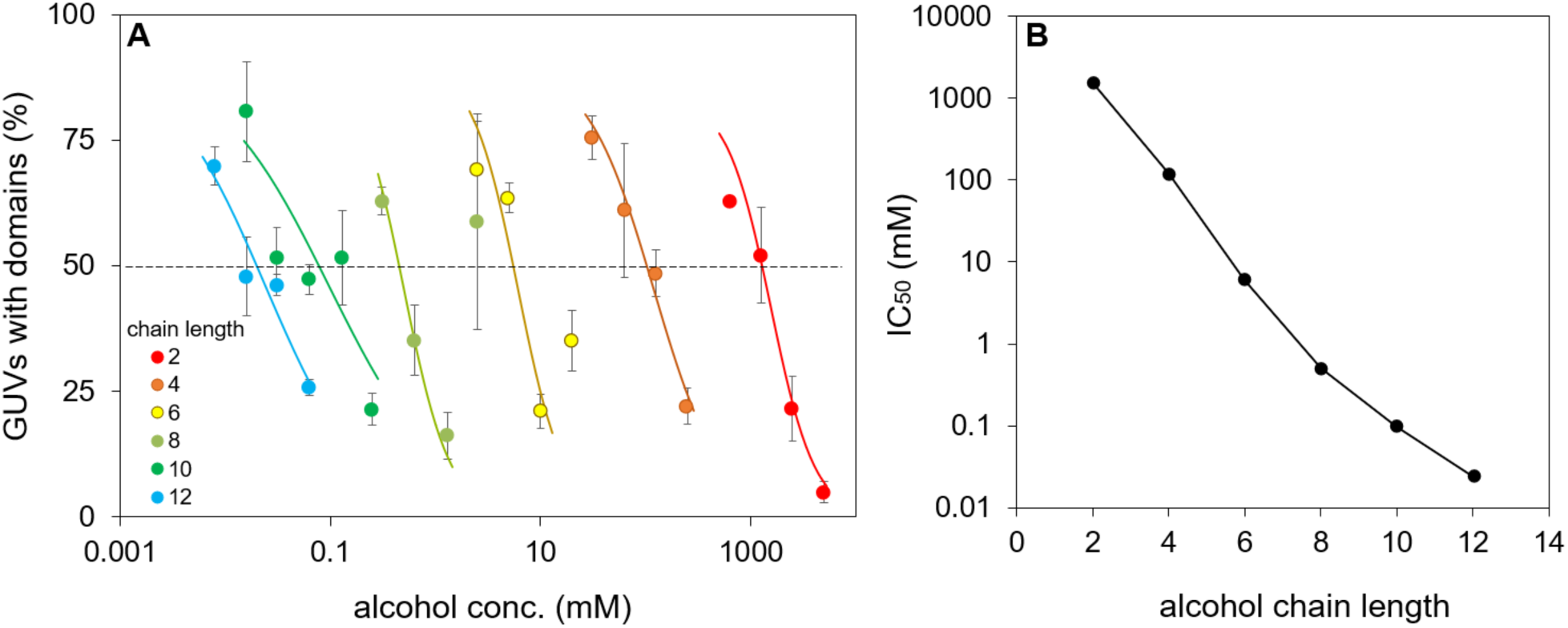
The relationships between IC50s of domain formation in GUVs and the chain lengths of alcohols. (*A*) The percentages of the GUVs with domains are plotted against the concentrations of the alcohols (C_2_-C_12_). The lipid composition of GUV was the same as that in yellow circle in fig. 1. Images were recorded by confocal microscopy and more than 100 GUVs were counted for each alcohol concentration in one experiment. The logistic curves for the indicated alcohols are fitted to determine their IC_50_ values and the averages are indicated by lines. Data are the means ± SEM (*n* = 3). (*B*) The IC_50_ values of the alcohols are plotted against the chain lengths.

### The calculated membrane concentrations inhibiting domain formation are constant irrespective of the chain length

In the biomembrane, the membrane-water partition coefficients (*K*_mw_: *C*_m_/*C*_w_) of alcohols increase exponentially with chain length, as do the octanol-water partition coefficients (*P*_ow_: *C*_o_/*C*_w_) (Table 1).^4^ Thus, the product of *K*_mw_ and the IC_50_ (*C*_w_) determined in GUV provide an estimate for the alcohol concentration in a pure membrane without proteins (*C*_m_) when alcohol exerts biological effects. In the GUVs, the difference among the IC_50_ values (C_2_−C_12_) was 62500-fold, whereas that among their *C*_m_ values was 2.8-fold (Table 1). The effective doses (ED_50_) of the alcohols (C_2_−C_12_) for anesthesia of tadpoles also decrease exponentially with chain length according to the MO correlation. The difference between the ED_50_ values (*C*_w_) was 22222-fold, whereas the difference between their C_m_ values calculated by the *K*_mw_ and ED_50_ values was 3.2-fold.^4^ In the case of MIC for yeast growth, the difference between the yeast MICs (C_2_−C_12_) was 39700-fold, whereas the difference between their C_m_ values calculated by the *K*_mw_ values and the MICs was 1.5-fold (Table 1).^13^ Collectively, the differences between the *C*_m_ values are small in GUVs and living cells, whereas the differences between the *C*_w_ values are large. Accordingly, the constant value in *C*_m_ results from the direct interaction of alcohols with membrane lipids that are common in GUVs and living cells but not with proteins. Based on these observations, alcohols would exert biological effects irrespective of their chain lengths after they reach a constant concentration in the membrane (even in the GUVs free from proteins), which would be the principle underlying the MO correlation.

## Discussion

Here, by monitoring the inhibition of domain formation in GUVs, we demonstrated that the MO correlation by alcohols was established in the lipid membrane without proteins. Inhibition was also observed in GPMVs treated with *n*-alcohols and the general anesthetic propofol^27^ and in GUVs treated with local anesthetics,^36,37^ although the MO correlation was not confirmed. Thus, the inhibition would not be caused by a specificity for alcohols but rather by their common amphiphilic structures that can penetrate within lipids. GUVs consisting of 1,2-dipalmitoyl-*sn*-glycero-phosphocholine (16:0-PC, DPPC)/DOPC/CHOL are often used to investigate the effect of anesthetics. However, the MO correlation has not been confirmed in the DPPC-GUV system because alcohols (C_10_ and C_14_) inhibit domain formation at the same temperature, whereas alcohols (C_2_-C_8_) stabilize domain formation.^35^ Therefore, this is the first report to reproduce the MO correlation in the lipid membrane free from proteins (fig. 3). Most likely, the discrepancy with previous reports resulted from the difference in lipid composition. RBL-2H3 cell-derived GPMVs contain more C_18:0_ than C_16:0_,^38^ and negatively charged lipids, such as phosphatidylserines and phosphatidyl inositols.^39^ Our GUVs contain DSPC (18:0-PC) instead of DPPC (16:0-PC) and negatively charged lipids, 18:0-PG and 18:1-PG. Thus, our GUVs reflect the properties of GPMV more than DPPC-GUV, although the detailed mechanism is unclear. These GUVs might be useful to effectively evaluate positively charged drugs, such as local anesthetics.

There are several mechanisms that lead to the inhibition of domain formation by alcohols. The addition of 1-palmitoyl-2-oleoyl-*sn*-glycero-phosphocholine (16:0,18:1-PC, POPC) to the DSPC/DOPC/CHOL system was reported to reduce the domain to the nanoscopic size.^40^ Similarly, alcohols might reduce the domain to a size undetected by ordinary fluorescence microscopy. Alternatively, the amphiphilic alcohols could penetrate nonspecifically within the phosholipids consisting of saturated DSPC and unsaturated DOPC according to their amphiphilic orientation. This nonspecific penetration might eliminate the biased interaction among the phospholipids and cholesterol, thereby resulting in the disappearance of phase separation for domain formation. Since similar disappearance was observed at high contents of amphiphilic cholesterol (fig. 1B), this effect may be common for chemicals with amphiphilic structures.

The variance among the membrane concentrations (*C*_m_) of the alcohols (C_2_-C_12_) inhibiting domain formation in GUVs was 2.8-fold; thus, *C*_m_ remains constant regardless of the chain length (Table 1). The constant value of *C*_m_ by the alcohols is also calculated by combining the *K*_mw_ values and various biological doses, such as the ED_50_ values obtained for tadpole anesthesia^4^ and anti-hemolysis of human erythrocytes,^41^ as well as the MICs (Table 1)^12^ and the LD_50_ values obtained for yeast,^13^ as observed in the *C*_m_ values calculated by other chemicals.^10^ Thus, the constant values of *C*_m_ in GUVs reflect the values in the biomembranes when alcohols exert various biological effects, although their absolute values of *C*_m_ vary depending on the subjects of research.

In comparison with *C*_m_, the variance among the octanol phase concentrations (*C*_o_) of the alcohols (C_2_-C_12_), which was calculated by the products of the IC_50_ values in GUVs and the octanol-water partition coefficients (*P*_ow_: *C*_o_/*C*_w_), was 7.2-fold. This variance was larger than the 2.8-fold among the *C*_m_ values (Table 1) The difference between the *C*_m_ and *C*_o_ values was also observed for MIC of yeast growth (*C*_m_’s: 1.5-fold, *C*_o_’s: 7.6-fold, Table 1), LD_50_ of yeast viability (*C*_m_’s: 2-fold, *C*_o_’s: 7-fold),^13^ and ED_50_ of tadpole anesthesia (*C*_m_’s: 3.2-fold, *C*_o_’s: 12.4-fold).^4^ Thus, the difference between the *C*_m_ and *C*_o_ values in GUVs reflects those observed in the biomembranes of yeast and tadpole. These differences are because the *C*_o_ values for C_10_ and C_12_ are larger than those for C_2_-C_8_, which is attributed to the *P*_ow_ values for C_10_ and C_12_ larger than the *K*_mw_ values (Table 1). The amphiphilic alcohols (C_2_-C_8_) would be readily solubilized in octanol (C_8_) and vesicles via penetration according to their amphiphilic orientations. The longer alcohols (C_10_-C_12_) would also be solubilized in the vesicles with long alkyl chains via penetration; however, they might be solubilized in octanol with a shorter chain at a high concentration via chemically stable formation of micelle or emulsion, resulting in high *P*_ow_. Presumably, this is responsible for the difference between *C*_m_ and *C*_o_. Taken together, these observations suggest that the constant value in *C*_m_ arises from the direct interaction between alcohols and highly lipophilic membrane lipids with long alkyl chains, rather than octanol with a restricted hydrophobicity.

In conclusion, when alcohols reach the same critical concentration in the lipid membrane (*C*_m_), similar biological effects appear irrespective of the chain length. To maintain the same concentration in the membrane, the apparent concentrations in the aqueous phase (*C*_w_) decrease exponentially with their lipophilicities (partition coefficients), thereby resulting in the MO correlation. In general, the protein theory focusing on *C*_w_ states that a highly lipophilic compound targets minor membrane proteins in a highly specific manner due to its effects at a low *C*_w_. In contrast, the lipid theory focusing on *C*_m_ states that the compound targets major or various membrane proteins because its actual *C*_m_ around the proteins would be high.

## Acknowledgments

The authors would like to thank Gerald W. Feigenson for supervising to prepare artificial membrane, Yi Wen and Thais Azevedo Enoki for technical help (Cornell University, Ithaca, New York, USA), and Ichiro Terashima for critical reading of the manuscript (The University of Tokyo, Tokyo, JAPAN). The authors acknowledge the financial support from the Sasakawa Scientific Research Grant (29-637 to AM), JSPS Overseas Challenge Program for Young Researchers (201880084 to AM), and JSPS KAKENHI (19J13923 to AM).

## References

1. Meyer H: Zur theorie der Alkoholnarkose. Arch Exp Pathol Pharmakol 1899; 42:109–18

2. Gill CO, Ratledge C: Toxicity of n-alkanes, *n*-Alk-1-enes, *n*-alkan-1-ols and *n*-alkyl-1-bromides towards yeasts. Microbiology 1972; 72:165–72

3. Teh JS: Toxicity of short-chain fatty acids and alcohols towards *Cladosporium* resinae. Appl Microbiol 1974; 28:840–4

4. Pringle MJ, Brown KB, Miller KW: Can the lipid theories of anesthesia account for the cutoff in anesthetic potency in homologous series of alcohols? Mol Pharmacol 1981; 19:49−55

5. Kubo I, Muroi H, Kubo A: Structural functions of antimicrobial long-chain alcohols and phenols. Bioorg Med Chem 1995; 3:873−80

6. Uesono Y, Toh-E A, Kikuchi Y, Terashima I: Structural analysis of compounds with actions similar to local anesthetics and antipsychotic phenothiazines in yeast. Yeast 2011; 28:391−404

7. Ingólfsson HI, Andersen OS: Alcohol’s effects on lipid bilayer properties. Biophys J 2011; 101:847−55

8. Hansch C: Quantitative approach to biochemical structure-activity relationships. Acc Chem Res 1969; 2:232−9

9. Antkowiak B: How do general anaesthetics work? Naturwissenschaften 2001; 88:201−13

10. Meyer KH, Hemmi H: Beitrage zur Theorie der Narkose, III. Biochem Z 1935; 277:39–71

11. Franks NP, Lieb WR: Molecular and cellular mechanisms of general anaesthesia. Nature 1994; 367:607–14

12. Matsumoto A, Uesono Y: Physicochemical Solubility of and Biological Sensitivity to Long-Chain Alcohols Determine the Cutoff Chain Length in Biological Activity. Mol Pharmacol 2018; 94:1312−20

13. Matsumoto A, Adachi H, Terashima I, Uesono Y: Escaping from the Cutoff Paradox by Accumulating Long-Chain Alcohols in the Cell Membrane. J Med Chem 2022; 65:10471–80

14. Pavel MA, Petersen EN, Wang H, Lerner RA, Hansen SB: Studies on the mechanism of general anesthesia. Proc Natl Acad Sci USA 2020; 117:13757–66

15. Pike LJ: Rafts defined: a report on the Keystone Symposium on Lipid Rafts and Cell Function. J Lipid Res 2006; 47:1597–8

16. Kingsley PB, Feigenson GW: The synthesis of a perdeuterated phospholipid: 1,2-dimyristoyl-*sn*-glycero-3-phosphocholine-d_72_. Chem Phys Lipids 1979; 24:135–47

17. Enoki TA, Heberle FA, Feigenson GW: FRET detects the size of nanodomains for coexisting liquid-disordered and liquid ordered phases. Biophys J 2018; 114:1921–35

18. Zhao J, Wu J, Heberle FA, Mills TT, Klawitter P, Huang G, Costanza G, Feigenson GW: Phase studies of model biomembranes: Complex behavior of DSPC/DOPC/Cholesterol. Biochim Biophys Acta Biomembr 2007; 1768:2764–76

19. Akashi K, Miyata H, Itoh H, Kinoshita JrK: Preparation of giant liposomes in physiological conditions and their characterization under an optical microscope. Biophys. J 1996; 71:3242–50

20. Schindelin J, Arganda-Carreras I, Frise E, Kaynig V, Longair M, Pietzsch T, Preibisch S, Rueden C, Saalfeld S, Schmid B, Tinevez J, White DJ, Hartenstein V, Eliceiri K, Tomancak P, Cardona A: Fiji: an open-source platform for biological-image analysis. Nat Methods 2012; 9: 676–82

21. Kemmer G, Keller S: Nonlinear least-squares data fitting in Excel spreadsheets. Nat Protoc 2010; 5:267–281

22. Ritz C: Toward a unified approach to dose-response modeling in ecotoxicology. Environ Toxicolo and Chem 2010; 29:220–229

23. Malinsky J, Opekarová M, Grossmann G, Tanner W: Membrane microdomains, rafts, and detergent-resistant membranes in plants and fungi. Ann Rev Plant Biol 2013; 64:501–29

24. Sáenz JP, Grosser D, Bradley AS, Lagny TJ, Lavrynenko O, Broda M, Simons K: Hopanoids as functional analogues of cholesterol in bacterial membranes. Proc Nat Acad Sci USA 2015; 112:11971–6

25. Sezgin E, Levental I, Mayor S, Eggeling C: The mystery of membrane organization: composition, regulation and roles of lipid rafts. Nat Rev Mol Cell Biol 2017; 17:361–74

26. Kamaya H, Matubayasi N, Ueda I: Biphasic effect of long-chain *n*-alkanols on the main phase transition of phospholipid vesicle membranes. J Phys Chem 1984; 88:797–800

27. Gray E, Karsklake J, Machta BB, Veatch SL: Liquid general anesthetics lower critical temperatures in plasma membrane vesicles. Biophys J 2013; 105:2751–9

28. van Meer G, Voelker DR, Feigenson GW: Membrane lipids: where they are and how they behave. Nat Rev Mol Cell Biol 2008; 9:112–24

29. Korlach J, Schwille P, Webb WW, Feigenson GW: Characterization of lipid bilayer phases by confocal microscopy and fluorescence correlation spectroscopy. Proc Nat Acad Sci USA 1999; 96:7563–8

30. Veatch SL, Keller SL: Organization in lipid membranes containing cholesterol. Phys Rev Lett 2002; 89:268101–4

31. Baumgart T, Hunt G, Farkas ER, Webb WW, Feigenson GW: Fluorescence probe partitioning between L_o_/L_d_ phases in lipid membranes. Biochim Biophys Acta Biomembr 2007; 1768:2182–94

32. Baumgart T, Hammond AT, Sengupta P, Hess ST, Holowka DA, Baird BA, Webb WW: Large-scale fluid/fluid phase separation of proteins and lipids in giant plasma membrane vesicles. Proc Nat Acad Sci USA 2007; 104:3165–70

33. Veatch SL, Cicuta P, Sengupta P, Honerkamp-Smith A, Holowka D, Baird B: Critical fluctuations in plasma membrane vesicles. ACS Chem Biol 2008; 3:287–93

34. Veatch SL, Keller SL: Miscibility phase diagrams of giant vesicles containing sphingomyelin. Phys Rev Lett 2005; 94:3–6

35. Cornell CE, McCarthy NLC, Levental KR, Levental I, Brooks NJ, Keller SL: *n*-Alcohol length governs shift in *L*_o_-*L*_d_ mixing temperatures in synthetic and cell-derived membranes. Biophys J 2017; 113:1200–11

36. Sugahara K, Shimokawa N, Takagi M: Destabilization of phase-separated structures in local anesthetic-containing model biomembranes. Chem Lett 2015; 44:1604–6

37. Kinoshita M, Chitose T, Matsumori N: Mechanism of local anesthetic-induced disruption of raft-like ordered membrane domains. Biochim Biophys Acta Gen Subj 2019; 1863:1381–9

38. Kawasaki M, Toyoda M, Teshima R, Sawada J, Saito Y: Effect of α-linolenic acid on the metabolism of ω-3 and ω-6 polyunsaturated fatty acids and histamine release in RBL-2H3 cells. Biol Pharm Bull 1994; 17:1321–5

39. Chang E, Zheng Y, Holowka D, Baird B: Alteration of lipid composition modulates FcεRI signaling in RBL-2H3 cells. Biochemistry 1995; 34:4376–84

40. Konyakhina TM, Wu J, Mastroianni JD, Heberle FA, Feigenson GW: Phase diagram of a 4-component lipid mixture: DSPC/DOPC/POPC/chol. Biochim Biophys Acta Biomembr 2013; 1828:2204–14

41. Roth S, Seeman P: The membrane concentrations of neutral and positive anesthetics (alcohols, chlorpromazine, morphine) fit the Meyer-Overton rule of anesthesia; negative narcotics do not. Biochim Biophys Acta 1972; 255:207−19

