## Supplemental Figures for "Establishment of the Meyer-Overton correlation in an artificial membrane without protein"

**Supplemental Figure S1. A schematic illustration of the gentle hydration method for GUV preparation.**

Lipid mixtures were dissolved with chloroform in a test tube (1) and dried into a thin film on a rotary evaporator at 60°C, followed by 1 h with high vacuum (2). The thin dry film was then hydrated by introducing wet $N_2$ gas at 60°C for 25 min. The prewarmed 100 mM glucose solution were added with the alcohols and incubated for 2 h at 60°C for GUV formation (3), followed by gently cooled over 10 h to 25°C (4). The GUVs were harvested into 100 mM sucrose solution containing the same concentration of the alcohols. The details are described in methods.

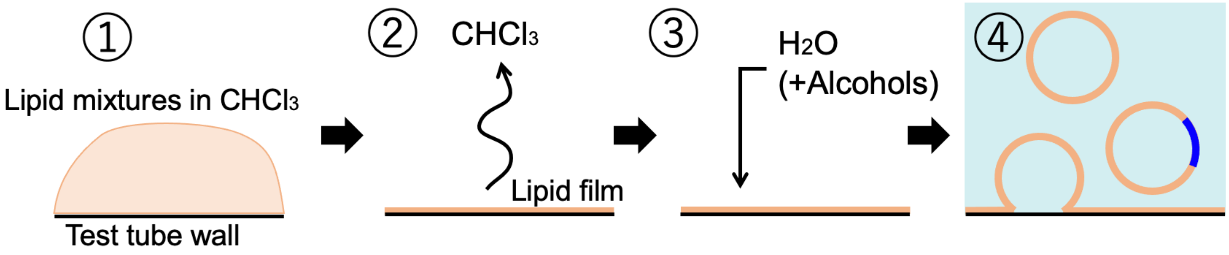

**Supplemental Figure S2. Ethanol did not affect the preformed membrane domains.**

The GUVs with the preformed domains (DSPC/DOPC/CHOL = 20/54/26, fig. 1A, yellow circle) were treated with 5 M of ethanol ($C_2$) for 15 min at 25°C. Bar, 20 μm. In contrast to the cases of domain formation in the presence of $C_2$ (fig. 2A and 2C), distinct domains were still observed in the GUVs.

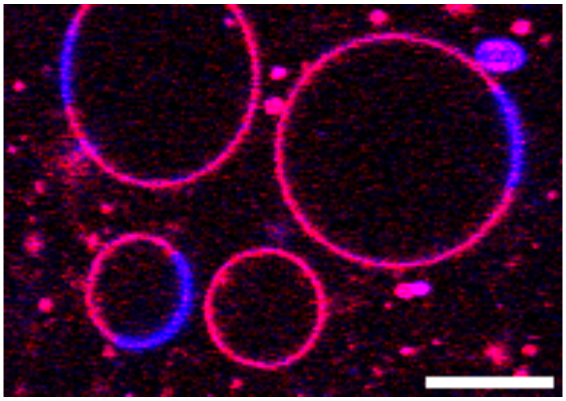
